# Using natural language processing to extract plant functional traits from unstructured text

**DOI:** 10.1101/2023.11.06.565787

**Authors:** Viktor Domazetoski, Holger Kreft, Helena Bestova, Philipp Wieder, Radoslav Koynov, Alireza Zarei, Patrick Weigelt

**Affiliations:** Biodiversity, Macroecology & Biogeography, University of Göttingen, Göttingen, Germany; Campus-Institute Data Science, Göttingen, Germany; Centre of Biodiversity and Sustainable Land Use, University of Göttingen, Göttingen, Germany; Gesellschaft für wissenschaftliche Datenverarbeitung mbH Göttingen, Göttingen, Germany

**Keywords:** Automatic Trait Extraction, Biodiversity, Functional Plant Ecology, Natural Language Processing, Large Language Models, Vascular Plants

## Abstract

Functional plant ecology aims to understand how functional traits govern the distribution of species along environmental gradients, the assembly of communities, and ecosystem functions and services. The rapid rise of functional plant ecology has been fostered by the mobilization and integration of global trait datasets, but significant knowledge gaps remain about the functional traits of the ∼380,000 vascular plant species worldwide. The acquisition of urgently needed information through field campaigns remains challenging, time-consuming and costly. An alternative and so far largely untapped resource for trait information is represented by texts in books, research articles and on the internet which can be mobilized by modern machine learning techniques.

Here, we propose a natural language processing (NLP) pipeline that automatically extracts trait information from an unstructured textual description of a species and provides a confidence score. To achieve this, we employ textual classification models for categorical traits and question answering models for numerical traits. We demonstrate the proposed pipeline on five categorical traits (growth form, life cycle, epiphytism, climbing habit and life form), and three numerical traits (plant height, leaf length, and leaf width). We evaluate the performance of our new NLP pipeline by comparing results obtained using different alternative modeling approaches ranging from a simple keyword search to large language models, on two extensive databases, each containing more than 50,000 species descriptions.

The final optimized pipeline utilized a transformer architecture to obtain a mean precision of 90.8% (range 81.6-97%) and a mean recall of 88.6% (77.4-97%) on the categorical traits, which is an average increase of 21.4% in precision and 57.4% in recall compared to a standard approach using regular expressions. The question answering model for numerical traits obtained a normalized mean absolute error of 10.3% averaged across all traits.

The NLP pipeline we propose has the potential to facilitate the digitalization and extraction of large amounts of plant functional trait information residing in scattered textual descriptions. Additionally, our study adds to an emerging body of NLP applications in an ecological context, opening up new opportunities for further research at the intersection of these fields.

## 1 Introduction

Functional plant ecology is a field that has been exponentially growing in the last few decades. With more than 380,000 vascular plants already identified around the globe, there is a necessity to understand these species functionally in order to analyze and answer fundamental questions about their role in maintaining ecosystem functioning (Antonelli, 2023). However, biases in and lack of trait data for many species and traits hamper solid inference on the spatial distribution of traits, their responses to the environment and their importance for ecological processes (Maitner et al., 2023).

Several key databases contain information on plant traits, such as the TRY initiative (Kattge et al., 2011, Kattge et al., 2020), the Global Inventory of Floras and Traits (GIFT) (Weigelt et al., 2020), the Botanical Information and Ecology Network database (BIEN) (Maitner et al., 2018), the Open Traits Network (Gallagher et al., 2019) and the Austraits database (Falster et al., 2021). These databases hold information on hundreds of thousands of plant species and thousands of traits, varying from individual measurements to species-level information and from physiological, chemical and genomic traits to whole plant and structural traits, making them essential resources for a large number of ecological fields and across scales (König et al., 2019). This has led to a vast number of studies in the field of functional ecology and tremendously enhanced our understanding of the global distribution of plant form and function (Díaz et al., 2016, Wright et al., 2004, Moles et al., 2007).

However, these landmark papers are based on only a small proportion of the plant species on Earth due to a lack of trait data. Although available trait databases contain a large amount of data, only a small fraction of described plants are covered, with global mean trait completeness of 0.21% and global median trait completeness of 0.0051% in the TRY database (Maitner et al., 2023, Kattge et al., 2020). Furthermore, the information contained is strongly biased in multiple dimensions (Maitner et al., 2023, Meyer et al., 2016, König et al., 2019). While information for traits such as growth form may be well represented, other, equally important traits such as specific leaf area may not. In addition, the availability of traits is taxonomically and spatially biased, depending on socioeconomic drivers (Maitner et al., 2023). While trait imputation based on trait-trait correlations and phylogenetic relationships among species may help to fill gaps (Schrodt et al., 2015), it only works for taxonomically and geographically representative data (Penone et al., 2014). Particularly for analyses at a fine spatial grain (Bruelheide et al., 2018), high trait resolution (i.e. many very specific traits), or analyses of undersampled regions (e.g. the tropics), the only way forward is the mobilization of more trait data.

New trait data may come from extensive field campaigns and ecological experiments. However, these approaches are time-consuming and costly, therefore, an additional promising and potentially faster and cheaper option is to mobilize available but so far untapped information hidden in the wealth of published and online literature. Regional floras, checklists, and taxonomic monographs, for example, contain descriptions of the species giving more or less detailed information on morphological, reproductive, dispersal, and other traits (Weigelt et al., 2020). In addition, a huge amount of information is contained in primary research articles (e.g. species descriptions) and many regional to global websites (e.g. Wikimedia, JSTOR Plant Science, Plants of the World Online). So far, such data had to be manually extracted page by page in search of specific trait information, but the emergence of powerful machine learning techniques opens up new avenues for extracting this information in an automated way.

The past decade has seen an exponential rise in the application of machine learning (ML) for a variety of tasks, including ecological problems. As data sources continue to grow, the ability to automatically learn patterns from data becomes increasingly critical. Deep learning (DL) models, which utilize large numbers of hidden layers and nonlinear activations (LeCun et al., 2015), sped up the progress of fields such as computer vision, digital signal processing, and natural language processing (NLP).

Ecologists have recently started using ML and DL (Christin et al., 2019, Pichler et al., 2023), particularly for tasks in wildlife conservation (Tuia et al., 2022) like animal detection in camera traps (Steenweg et al., 2017) and species and individual recognition in bioacoustic signals (Sugai et al., 2019). By coupling images from iNaturalist and traits from the TRY database, convolutional neural networks (CNN) have been used to predict trait values (Schiller et al. 2021). Similarly, CNNs have been used CNNs to measure functional traits of skeletal museum specimens (Weeks et al. 2022).

NLP algorithms may be applied to numerous scientific ends, however, one of the most important may be to extract information from unstructured texts (Singh, 2018). Despite the success of DL models in ecology using tabular, image, and acoustic data, so far, few studies have explored the potential of NLP models for ecological data analysis and interpretation and ecologists and evolutionary biologists have been slower to adopt NLP models in comparison to other fields, such as biomedicine or economics (Farell et al., 2022). So far, most modeling approaches have been applied to detect trends and topics and perform evidence synthesis and literature reviews (Farell et al., 2022). For example, NLP has been used to classify whether literature is relevant (Cornford et al., 2021) to the Living Planet Database (LPD: http://livingplanetindex.org/data_portal) and the PREDICTS database (Hudson et al., 2014). Furthermore, NLP models offer a unique opportunity to extract and integrate ecological data from unstructured text. One such application is TaxoNERD (Le Guillarme & Thullier, 2021), a tool that uses named entity recognition DL models to automatically extract taxonomic name information from ecological documents. Similarly, NLP has previously been used to identify and extract functional trait information (Endara et al., 2018, Mora & Araya, 2018). However, most of these approaches have stuck to simpler models such as dictionaries, term co-occurrences, or bag of words models, which have disadvantages in terms of complexity and predictive power.

Here, we propose to use NLP techniques to automatically extract functional trait data from unstructured texts. First, we provide a brief overview of the pipeline and formulate the problem of trait value prediction of categorical and numerical traits as two standard NLP tasks: classification and question answering. Furthermore, we elaborate on the process of data acquisition, preprocessing, and model selection. Using two sources of textual species descriptions, we train and evaluate a range of models, from a straightforward keyword search to state-of-the-art large language models^1^. Finally, we underline the challenges and promise in the current state of the field and offer ideas on how to proceed.

## 2 Materials & methods

### 2.1 Task formulation

In the proposed pipeline to extract functional traits from text, we start with a textual description of a species. This description can be taken from a variety of sources such as floras, scientific papers, and plant databases and usually contains species trait information for a few to dozen traits. Following standard NLP preprocessing, it is then used as the input for ML or DL models. The output of the models is a predicted trait value with a corresponding confidence score. We use two different supervised NLP tasks for the categorical and numerical traits respectively. The pipeline (Fig. 1.) is described in depth in the sections that follow.

**Figure 1:**
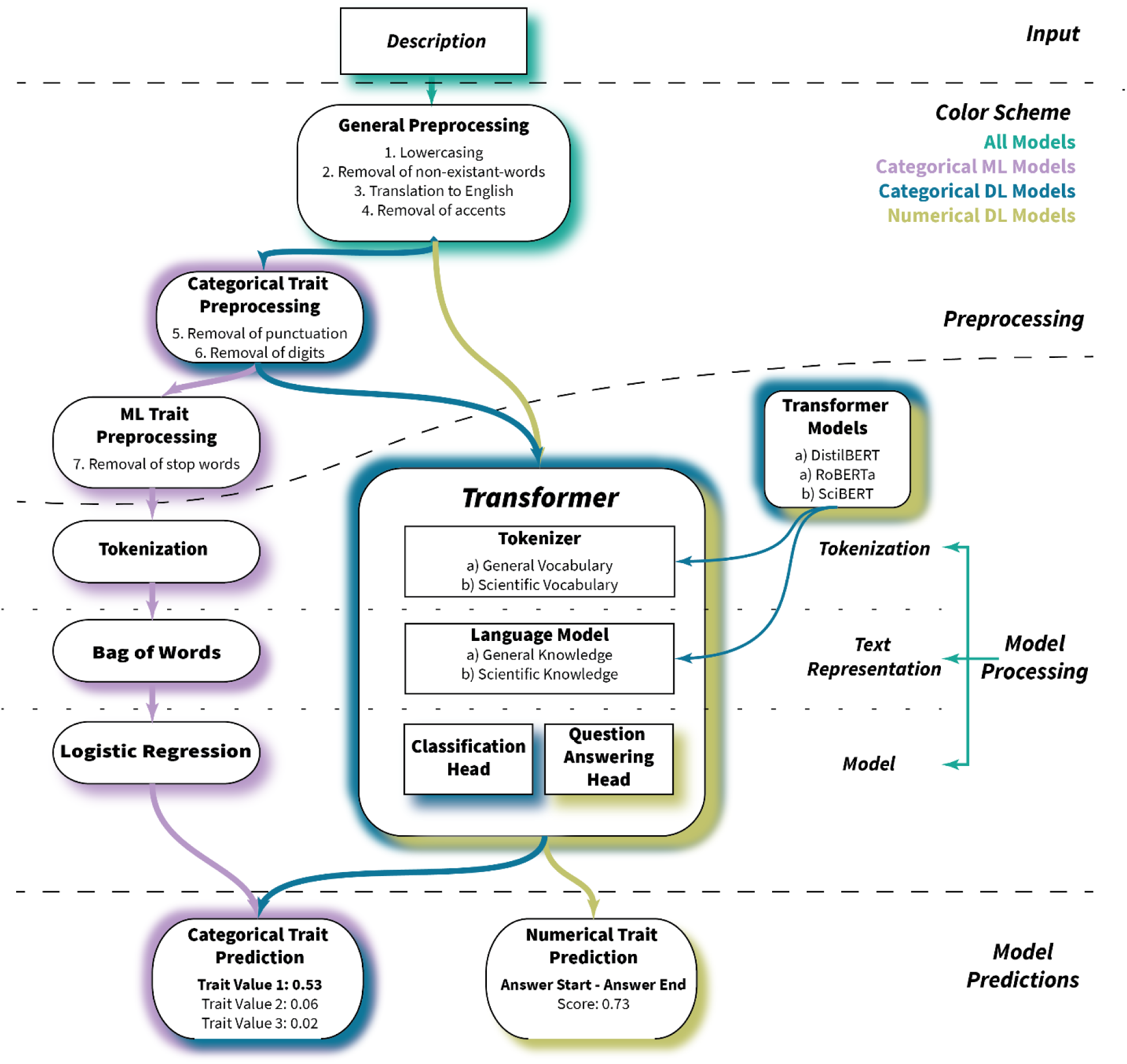
Proposed NLP pipeline used for the prediction of categorical and numerical traits based on textual species descriptions. The description first goes through a general preprocessing pipeline and then enters the categorical machine learning (ML) pipeline (violet), categorical deep learning (DL) pipeline (blue), and numerical DL pipeline (green). Within the corresponding pipelines, the description is further preprocessed if necessary and then tokenized and transformed into a vector embedding. Finally, the vector embedding is input in the corresponding model head and the model returns a prediction with an associated confidence score. An example description being processed using the pipeline is presented in Fig. S1.

#### 2.1.1 Categorical traits - sequence classification

Categorical traits are traits that have a discrete number of possible values. We considered five such traits, as defined in the GIFT database (Weigelt et al., 2020), representing important structural and life history aspects of plants (Taylor et al., 2023). We slightly modified some trait definitions, e.g. removed the biennial lifecycle, due to the very low representation in the data (less than 1%). Including the biennial lifecycle would have resulted in a very imbalanced dataset which would have to be approached with a different set of tools like few-shot learning (Wang et al., 2020).

● **Growth form**: herb, shrub, tree *(combination possible)*
● **Epiphyte**: epiphyte, terrestrial
● **Climber**: climber, self-supporting
● **Lifecycle**: annual, perennial *(combination possible)*
● **Life form**: phanerophyte, chamaephyte, hemicryptophyte, cryptophyte, therophyte *(combination possible)*

Given the finite number of classes per trait, predicting a categorical trait value can be considered a sequence classification task. Unlike keyword search algorithms, which rely on manually defined rules to predict the trait value, ML and DL models may automatically assign weights to words or phrases that are relevant to the trait value. For each trait, we trained a multi-class classification model, from which we get a probability for each trait value. Using these probabilities, there are two ways to get a prediction. The first method, which we used as the default for our models, is to pick the trait value with the highest probability. However, this approach has a few limitations: it can only predict one trait value, which precludes the use of combinations and it may provide a prediction even if the probability of all traits is low. The second way to obtain the prediction is to predict all trait values with a probability above a defined threshold t. This allows for combinations of trait values such as herb/shrub if the probabilities of both traits are above the threshold. Additionally, if there is insufficient support for a trait, the probabilities should be below the threshold and therefore no value will be predicted.

#### 2.1.2 Numerical traits – extractive question answering

Numerical traits are those described by a continuous numeric variable. In this paper, we considered three such traits that are commonly reported in species descriptions and encapsulate important information on plant functional strategy (Díaz et al., 2016, Wright et al., 2004):

● **Maximum plant height**, expressed in *m*
● **Maximum leaf length**, expressed in *cm*
● **Maximum leaf width**, expressed in *cm*

When predicting categorical traits, the occurrence of individual words can have a significant impact. For example, the word “annual” in the description increases the likelihood of the growth form herb since by definition only herbs are annual plants. In numerical traits, statistical inference of this kind is impossible. This led us to adopt an extractive, or contextual, question answering (QA) model to constrain the prediction within the information of the text. Extractive QA models ask a question (e.g. What is the height of the plant?) and return an answer and confidence score based on a context paragraph, in our case, the species description. Due to the nature of the model, it also returns the unit along with the numerical value, making it simple to convert between units. To get the final predicted number, we post-processed the answer of the QA model such that all answers that didn’t contain a number or a unit were filtered out. Furthermore, we filtered out all predictions with a confidence score below a defined threshold *t*. If the answer contained two numbers, we extracted the second value, under the assumption that they represented a range (minimum-maximum). Lastly, this value was transformed into the required unit of measurement for the trait.

### 2.2 Data

All of the above-defined problems are supervised NLP tasks and thus require textual data and corresponding labels. For this reason, we performed a web scrape to acquire species descriptions from two large online plant knowledge bases:

- Plants of the World Online (POWO), which aggregates information from regional floras.
- English Wikipedia Plant Articles (WIKI), which aggregates information written by volunteers.

We divided the descriptions from these databases by species and by source and then used them as the textual input in the NLP models. On the other hand, for the trait values that we used to train and evaluate the models, we extracted trait information from the Global Inventory of Floras and Traits (GIFT) database.

Data from Plants of the World Online (http://www.plantsoftheworldonline.org/) includes information on the taxonomy, and most importantly for us, textual descriptions of traits, identification, and distribution information per species. The descriptions are categorized hierarchically, containing information on various aspects such as leaf morphology, plant habit and reproductive information. We used the taxize R package (Chamberlain et al., 2020) to get the species description from the website, resulting in 288,254 descriptions for 59,151 plant species in 251 distinct categories. Using only descriptions of a category that contains information on a particular trait may improve model performance compared to using additional less relevant data that may lead to erroneous correlations. To test this hypothesis, we created one dataset using the entire Plants of the World Online knowledge base and two trait-specific datasets that use only the descriptions from certain categories. The POWO dataset was built by combining all descriptions per species per source and is the main one we used throughout the paper. The POWO_MGH used only descriptions from the Morphology General Habit category and contains relevant information for the traits: growth form, epiphyte, climber, lifecycle and plant height. Similarly, the POWO_ML dataset solely uses the Morphology Leaf category and includes data on leaf length and width. Further information on the Plants of the World dataset can be found in Supp. Information S2.1.

To get species descriptions from Wikipedia, we searched for English articles for all of the ∼270,000 species for which we found information for at least one of the selected functional traits in the GIFT database. We chose this direction because to train and evaluate the model, we need both textual descriptions and labels for each species. We used the Python Wikipedia-API (Wikipedia-API), and the Requests and Beautiful Soup (beautifulsoup4) web scraping libraries. This resulted in 194,994 descriptions for 55,631 species with description categories based on the sections in Wikipedia. Further information on the Wikipedia dataset can be found in Supp. Information S2.2.

The GIFT database (Weigelt et al. 2020) is a global archive of regional plant checklists and floras, and plant functional traits. It contains information for 109 traits for more than 290,000 species. Using the GIFT R-package v. 3.0 (Denelle et al. 2023), we extracted the values for the traits of interest and used them as labels in our models. It should be emphasized that not all traits have the same coverage in GIFT which leads to significant differences in the number of labeled samples per trait (Supp. Information S2, Tables S1, S2).

### 2.3 Data preprocessing

We used three different preprocessing pipelines on the descriptions of the POWO and WIKI datasets using the NLTK python library (Loper & Bird, 2002), resulting in descriptions for the categorical ML models, categorical DL models, and numerical DL models. We did this tailored to the different input requirements of the models. The base preprocessing pipeline consisted of the removal of artifacts and accents from the text. Using a keyword search that focused on language-specific terms, we detected non-English descriptions in the POWO dataset. We found 20,034 non-English descriptions out of the total 288,254 descriptions, a large majority of which (19,945) were Spanish. These descriptions were then translated using the GoogleTranslate Python API (googletrans). The text was lowercased and split into tokens, which in our case may represent words, numbers, or punctuation. For the categorical models, we removed all digits and punctuation as they are not informative for this type of analysis. Finally, since English stop words like ”the” do not give any trait information, we removed them from the categorical ML model descriptions.

### 2.4 Models

#### 2.4.1 Keyword search

We compared the performance of the categorical models to a simple keyword search, a commonly used technique in the automatized extraction of traits (Coleman et al., 2023). To do this, we created a dictionary relevant to each trait and used a script based on regular expressions to classify the descriptions. The keywords were generally the name of the class (trait value) and any synonyms that might be found in the description. A full list of the keywords used can be found in the Supp. Information S3.

#### 2.4.2 Logistic regression

For the categorical ML modeling approach, we chose logistic regression, a parametric predictive classification model widely used in ecological research. The model assumes a linear relationship between the independent variables and the target variable and predicts a probability for each class. However, as the model requires numeric input, we first had to transform the textual input to a numeric vector space to create a text representation. To do this we use a technique called bag of words (BOW), commonly known as one-hot encoding. BOW first takes the most common words (here the top 1000) in the entire textual corpus to form the vocabulary of the model. Each description is then transformed into a vector of size 1000, whose value corresponds to the number of times the associated term appears in the description. The BOW representation vector is then used as the predictor in the logistic regression model, while the trait value is the outcome. We implemented the BOW and logistic regression model using the scikit-learn ML library (Pedregosa et al., 2011; Supp. Information S4).

#### 2.4.3 Transformers

Transformers (Vaswani et al., 2017) are large language models that transformed the field of NLP, and subsequently went on to transform other fields like computer vision and digital signal processing, sparking a revolution in how DL models are built. The notion of transfer learning, which states that knowledge acquired through one task or dataset may be applied to another in a similar context, is the driving force underlying transformers. The addition of the attention mechanism enables the model to learn long-range dependencies without the usage of recurrent layers like earlier models. This was the primary component that allowed the model to achieve state-of-the-art performance on a variety of NLP tasks.

Its architecture can consist of one or both of the following parts: an encoder and a decoder. Encoder models, like Google’s BERT (Devlin et al., 2018), are characterized as having “bi-directional” attention since at each stage the attention layers can access all words in the initial sentence. This allows them to excel at tasks where an understanding of the entire textual input is required, such as sequence classification and extractive QA, making them the go-to architecture for our problem. Decoder models, like OpenAI’s GPT (Radford et al., 2019, Brown et al., 2020), used as the basis of ChatGPT, are also called auto-regressive, as the attention layers can only access words before the current position, making them best suited for tasks involving text generation. Finally, encoder-decoder models like T5 (Raffel et al., 2023) use both of these modules and are primarily used in tasks that need to generate text dependent on an input text, such as translation and summarization. The three core components of a transformer are the tokenizer, the language model, and the task-specific head. The tokenizer learns the vocabulary of the model and converts the input text into machine-readable data. Tokenizers can generally be split into word-based, character-based, and subword-based. In our model, we used the WordPiece tokenizer (Sennrich et al., 2015) which is a subword tokenization algorithm. It is based on the idea that while frequently used words should be learned as a whole, uncommon terms should be decomposed into meaningful subwords. The language model is the most important component of the transformer as it includes the attention heads. It is trained through the objectives of masked language modeling and next-sentence prediction. By being able to predict the masked words within a sentence or predict the next sentence, the model learns the meaning of words and how they co-occur. As a result, words with some connection such as “Japan” and “sushi” are closer together in the embedding space compared to “Japan” and “pizza” (Mikolov et al., 2013). The task-specific head, which is layered on top of the pre-trained language model, is the last part of the architecture. In a procedure known as fine-tuning, the weights of the language model are frozen and the head is trained using the task-specific data. Fine-tuning is a common method in transfer learning that enables the model to learn task-specific features while retaining the general knowledge acquired during pre-training. As previously mentioned, we use a sequence classification head for categorical traits and an extractive QA head for the numerical traits.

We trained and evaluated the following three models using the Huggingface’s transformers Python library (Wolf et al., 2019; Supp. Information S4).

● DistilBERT: The DistilBERT model (Sanh et al., 2019) is a transformer model that is trained using a self-supervised fashion using the BERT model as a teacher, a process called knowledge distillation. This allows the model to achieve comparable results to BERT with less than 20% of the parameters (66 million compared to the 340 million of BERT), making the model much easier to train and use. More specifically, it is pre-trained using masked language modeling, where a part of a sentence is masked out and the model is trained to predict it. Furthermore, it is pre-trained with a distillation loss and cosine embedding loss, such that the prediction probabilities and hidden states of the model are as close to those of BERTs as possible. The texts that this model was trained on are the same as BERT, a large corpus comprising the Toronto Book Corpus and the English Wikipedia Corpus. Therefore, the vocabulary and language model of the original pre-trained DistilBERT are general knowledge related. To use this model in our categorical trait pipeline, we attached a sequence classification head and fine-tuned it using our species descriptions and trait data. For our numerical trait pipeline, we attached a QA head which has been fine-tuned on the SQuAD v1.1 dataset (Rajpurkar et al., 2016) which contains more than 100,000 question-answer pairs in a variety of contexts.
● SciBERT: SciBERT (Beltagy et al., 2019) uses the same architecture as BERT but is trained on papers from the corpus of semanticscholar.org with a size of 1.14M papers and 3.1B tokens. Consequently, SciBERT has its vocabulary (scivocab) that is built to best match the training corpus, meaning that the model consists of a more scientific knowledge language model and vocabulary. We used this model only for the categorical traits by fine-tuning it identically to DistilBERT.
● RoBERTa: The RoBERTa model (Liu et al. 2019) optimizes BERT’s pre-training process using only the masked language modeling objective. Within our study, for the extraction of numerical traits, we used a version of RoBERTa with a question answering head. The QA head had been fine-tuned on the SQuAD v2.0 dataset (Rajpurkar et al., 2018) which builds on the first version of the dataset by adding unanswerable questions. This should theoretically improve the model in identifying cases where no numerical traits exist in the description.

### 2.5 Evaluation

We split the textual descriptions and trait data into a training set (75% of the data) and a test set (25%). Because of the large amount of data, we only used one split instead of a cross-validation approach. We evaluated the categorical traits on the following classification metrics which are most commonly used to quantify the performance of classification models:

● **Accuracy** - The number of descriptions correctly classified into a trait value scaled by the total number of descriptions.

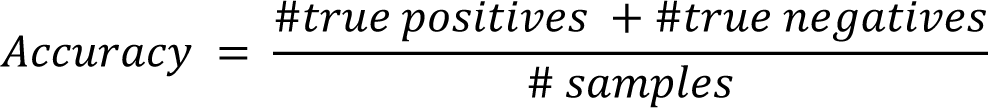

● **Precision** – The proportion of descriptions correctly classified into a given trait value scaled by the total number of descriptions classified into a given trait value.

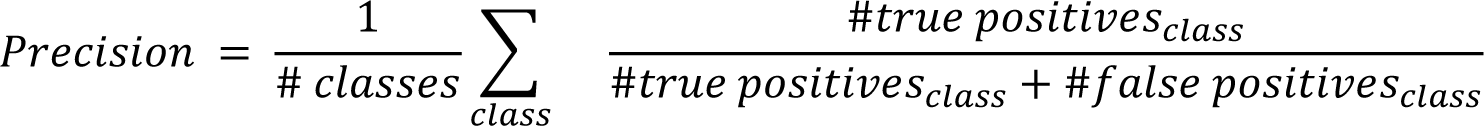

● **Recall** - The proportion of descriptions correctly classified into a given trait value scaled by the total number of descriptions that are labeled as a given trait value in the dataset.

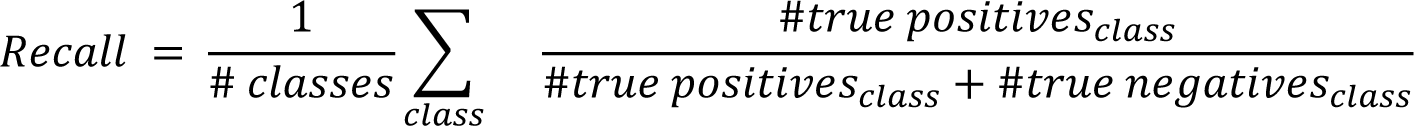

● **F1 score** - The harmonic mean of Precision and Recall.

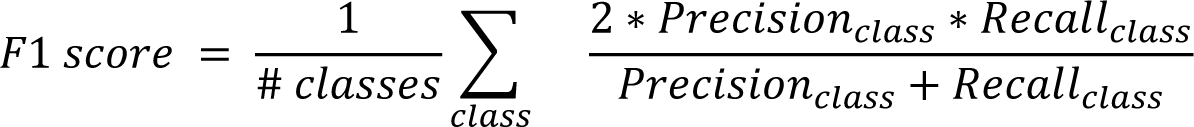

While accuracy is the most popular metric, it can be unrealistically high in imbalanced datasets such as ours, where the number of descriptions per trait value isn’t uniformly distributed. Therefore, we primarily focused on the remaining metrics. Since precision, recall, and f1 score are calculated for each class separately, we worked with the macro-version of the metric, meaning that we averaged over all trait values per trait. To evaluate how well the models performed on datasets with distinct characteristics from those used for training, we carried out an inter-dataset evaluation, where models trained using one dataset’s training set were evaluated on the test sets of the remaining two datasets (Supp. Information S5). Furthermore, when using the probabilistic output of the models combined with a threshold t as outlined in section 2.1.1, users can further customize the output. Since the probability threshold can be increased or decreased, users can get results with higher precision or a higher recall based on their needs. To analyze the model behavior dependent on the probability threshold, we calculated receiver operating characteristic and precision-recall curves, which show the precision versus recall at a variety of given thresholds. Furthermore, we analyze the model performance when using a threshold of t=0.5 and t=0.8 and compare it to our initial results. (Supp. Information S6).

To evaluate the numerical trait models, we first log-transformed the values due to their skewed distributions. To allow for the comparison of the results between the different datasets and models, the data was normalized to a range between 0 and 1, and we used the normalized mean absolute error (nMAE) as the main metric of model performance. In addition to this, we calculate the coverage, which represents the proportion of answers over the threshold that resulted in a trait value and unit of measurement. We separately evaluated the models on the aggregate POWO dataset compared to the trait-specific POWO_MGH, and POWO_ML datasets to evaluate the impact of the smaller data compared to the entire descriptions which might have a lot of interfering information. The threshold of the numerical models was set to *t=0.5* for DistilBERT and to *t=0.25* for RoBERTa. We used a lower threshold for RoBERTa as it is trained on the SQuAD v2.0 dataset which has unanswerable questions, thus the model should predict more skewed towards a score of 0 for descriptions with no trait information.

## 3 Results

### 3.1 Categorical traits

#### 3.1.1 Model performance comparison

The transformer models outperformed the standard keyword search model and the logistic regression model. On the POWO dataset, DistilBERT achieved an average precision across all traits of 90.6% and a recall of 88.5%. In comparison, the logistic regression model achieved an average precision of 88.4% and a recall of 85.1%, while the keyword search model had an average precision of 60.6% and an average recall of 37.7% (Fig. 2). The performance of SciBERT was comparable to that of DistilBERT, with an average improvement of 0.2% across metrics. This could be attributed to the scientific vocabulary and language model in SciBERT, which is relevant given that the POWO dataset includes a large amount of technical terminology. Alternatively, the increase in performance may be due to the higher number of parameters in the model.

**Figure 2:**
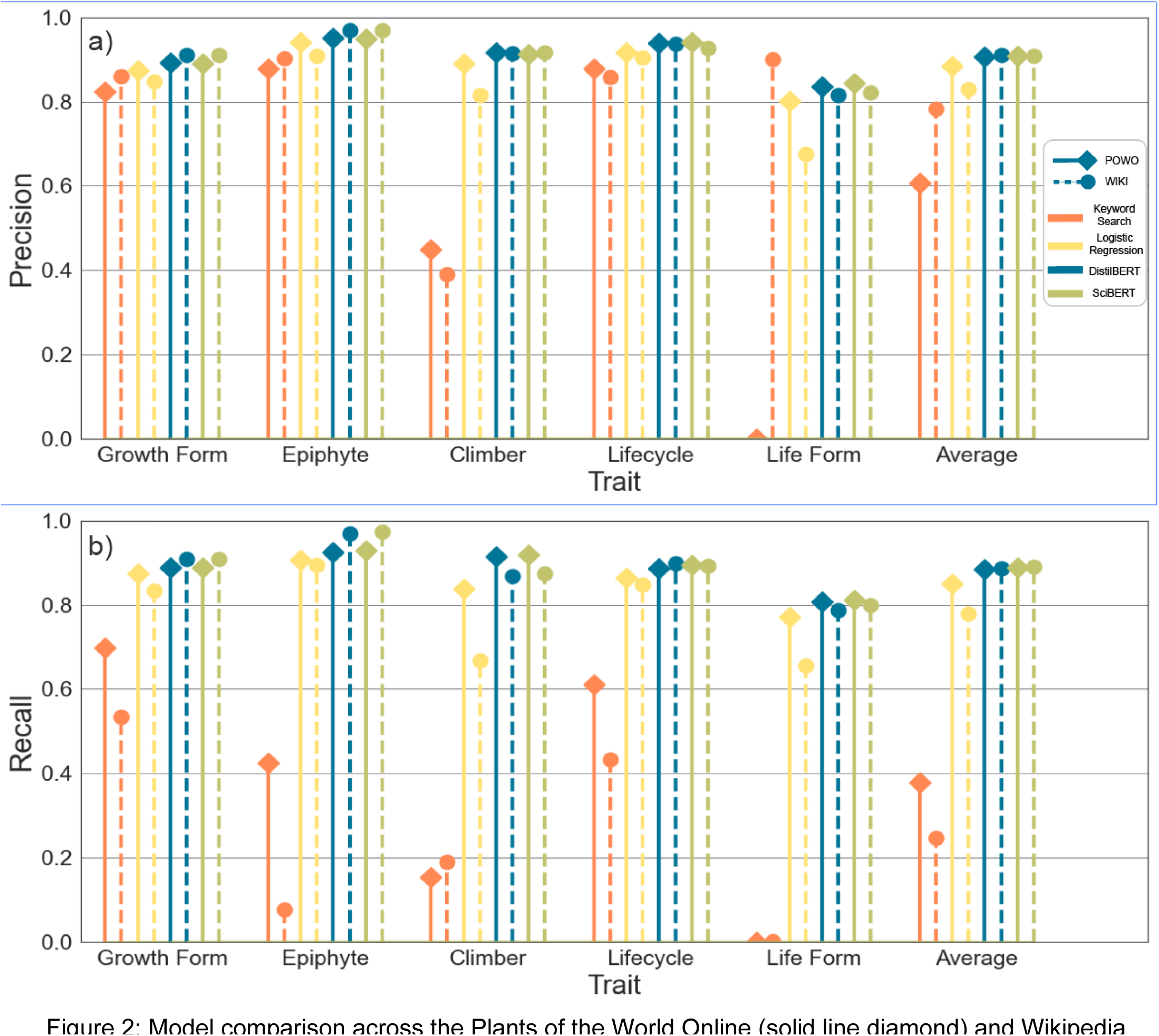
Model comparison across the Plants of the World Online (solid line diamond) and Wikipedia dataset (dashed line circle). Precision (a) and recall (b) for the categorical traits are shown for the keyword search (orange), logistic regression (yellow), DistilBERT (blue) and SciBERT (green).

**Fig. 3.**
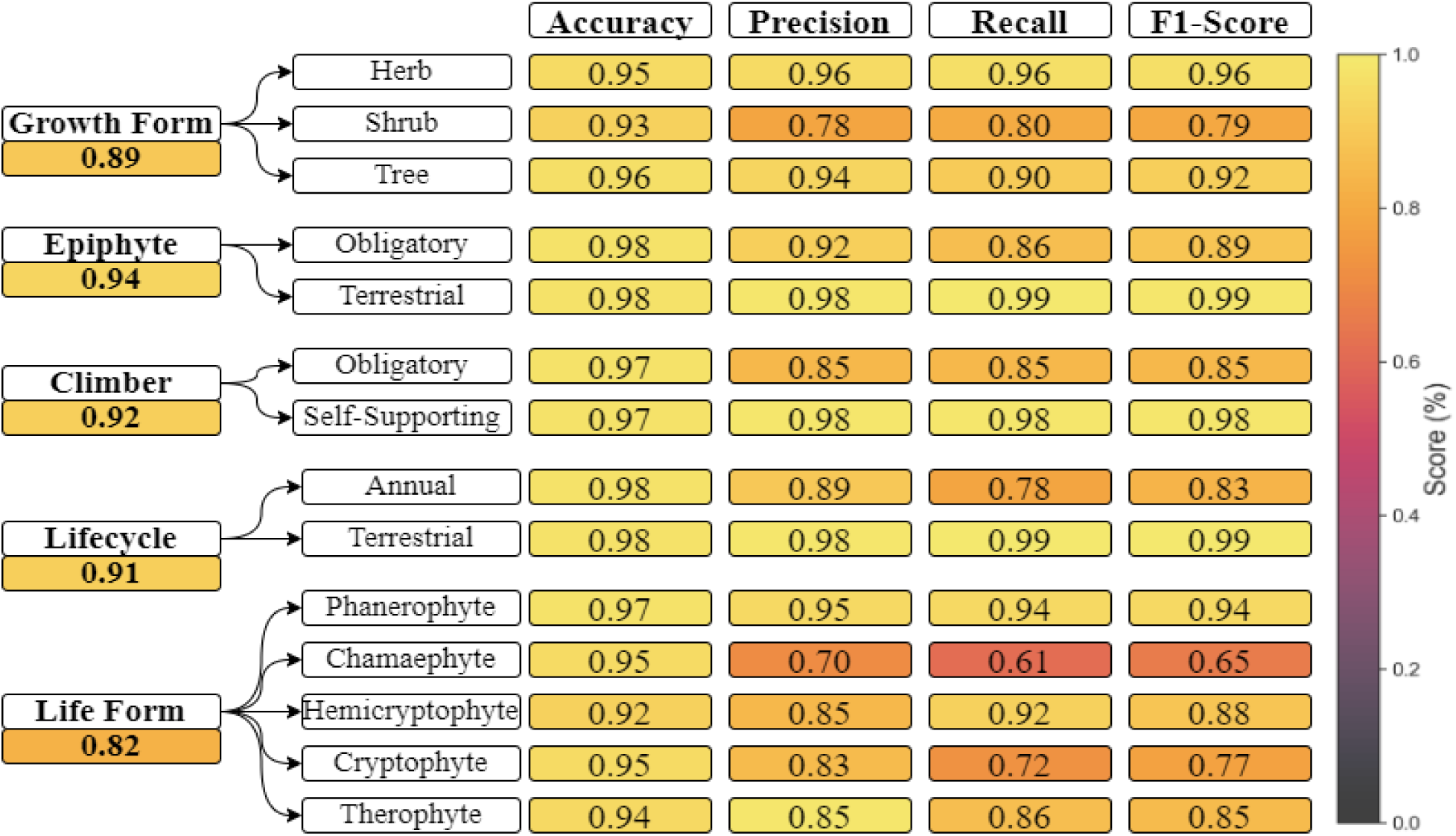
Class-specific performance metrics for each categorical trait value on the Plants of the World Online dataset using the DistilBERT model. The macro f1-score of the trait is shown below the trait name.

On the WIKI dataset, the keyword search had significantly lower predictions, with a recall of 24.7%, meaning that a large amount of these descriptions didn’t contain information on the trait. Similarly, the performance of the logistic regression model decreased by an average of 5% compared with the POWO dataset, reporting a precision of 83% and recall of 80%. Both of these techniques make trait predictions based on the occurrence of independent keywords, either defined by experts or calculated directly from the data, rather than considering the broader context. As the WIKI dataset contains a much smaller amount of expert information which directly signifies the traits, the decrease in performance of these models was expected. The transformers’ performance wasn’t affected by this factor and both models had a small increase in scores, with a precision of 91% and recall of 88.7% for the DistilBERT model, and a precision of 91% and recall of 89% for the SciBERT model.

#### 3.1.2 Performance variation between traits

DistilBERT and SciBERT showed the best results, however as DistilBERT had a much smaller number of parameters, we decided to focus on it for the rest of the analyses. The model performed best on the epiphyte trait with an f1 score of 93.8%/97%, and the worst on the life form trait with an f1 score of 82%/79.7% on the POWO/WIKI databases respectively. These differences can be explained by a few factors. One is whether the data is frequently included in the descriptions. While traits such as growth form are commonly reported in the descriptions we use, either explicitly or implicitly, as we can see in the recall rate of the keyword search, other traits like life form had less than 1% coverage in both datasets and a small number of implicit references making them much more ambiguous to detect. A second factor is the difficulty of the trait assessment. While some trait values can be determined based on simpler rules that the models can discover (e.g. annual -> herb), other trait values such as cryptophyte need either explicit or a large amount of implicit information to be classified. The final factor that may influence the results is the variability of the trait itself within the literature. While some traits like epiphyte and lifecycle have rather consistent values across sources, the boundaries between growth forms like herb and shrub or shrub and tree aren’t as clearly defined and are dependent on the source.

#### 3.1.3 Performance variation among traits

Due to the reasons outlined above, there were also big differences in model performance between trait values (Fig. 4). To explore these differences in more detail we analyzed class-specific metrics on the POWO dataset using the DistilBERT model. One trait value for which there was a decrease in performance is the growth form shrub which had approximately a 15% decrease in f1 score compared to the other two growth forms. The reason for this was the third factor explained above, as there often is no clear delineation between a herb and a shrub (e.g. subshrubs or semi-woody herbs) and even less so between a shrub and a tree. In some descriptions, a species is described as a herb while in the GIFT data, it is labeled as a shrub. Therefore, the variations in model performance could largely be attributed to the discrepancy between the description and label data, rather than the model’s inference capabilities. The trait for which we had the largest discrepancies, life form, suffered from the first two problems, the difficulty of assessment, and the lack of information in the description. For some trait values like phanerophyte, which has an f1 score of 94%, this wasn’t a problem as the model can discover the relation tree->phanerophyte, but for the other values like chamaephyte with an f1 score of 65%, this led to a significant decrease in the scores.

**Fig. 4.**
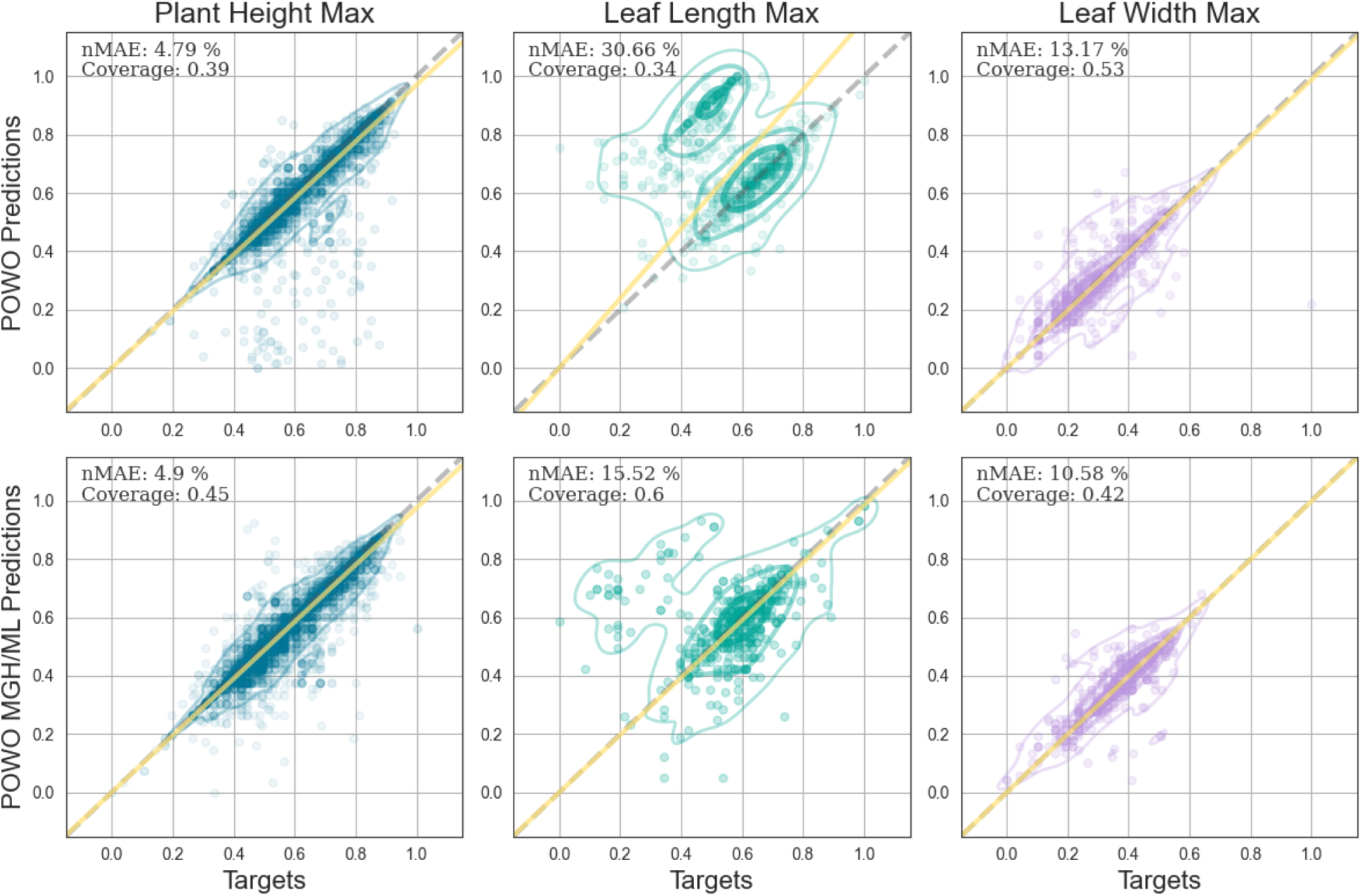
Observed vs. predicted numerical traits for the RoBERTa model on the aggregated Plants of the World Online and trait-specific Plants of the World Online datasets. The numerical traits are represented as plant height (blue), leaf length (cyan) and leaf width (violet). The 95% and 50% kernel density estimates are also shown by the corresponding trait color. The 1:1 line (gray dashed) and the regression line between the observed and predicted values (yellow solid) are also shown.

### 3.2 Numerical traits

Depending on the trait of interest and the database in question, 82 to 100% of the original answers contained a numeric value and a unit of measurement and were then used as input in the numerical models. The predictions on the trait-specific POWO datasets resulted in higher coverage of answers (50.83%) compared to the entire POWO dataset (39.17%). This could be because more descriptions of the entire POWO dataset do not contain information on the trait of interest. Consequently, the nMAE across all traits was also significantly decreased in the trait-specific datasets, with an average decrease of 5.74% for RoBERTa and 7.32% for DistilBERT.

RoBERTa achieved the smallest average nMAE of 10.64% across all traits on the trait-specific datasets. However, this varied significantly between traits, with a value of 5.3% for plant height and 16.12% for leaf length. This pattern was consistent across models and datasets, with the lowest errors for plant height, averaging 5.68%, and the highest for leaf length, averaging 24.39%. DistilBERT performed slightly worse than RoBERTa with an average nMAE of 13.1% on the trait-specific datasets. The coverage of this model was higher by 11% than RoBERTa, despite the smaller threshold of RoBERTa. However, as we are using descriptions and trait data from different sources, we do not know whether this increase is due to the higher recall of the model or due to false positive extractions. On top of this, the more compact architecture of DistilBERT led to a several-fold acceleration in the QA process which might be relevant for some tasks.

## 4. Discussion

The NLP workflow we proposed here has the potential to streamline the digitization and extraction of plant functional traits from textual descriptions. Consequently, an enormous amount of information currently hidden in unstructured texts online, in regional floras and in other scientific publications can be made digitally accessible for functional plant ecology studies. Manual trait extraction for the 8 traits of the 114.782 descriptions of the POWO and WIKI dataset, at a rate of 30 seconds per trait would take approximately 4.3 years of 40 hours per week work days. In contrast, the proposed workflow would extract the traits in a couple of hours without human supervision, saving time and resources. Overall, the categorical pipeline using DistilBERT achieved an average precision of 90.6%/91% and an average recall of 88.5%/88.7% across all traits on the POWO/WIKI datasets, a significant improvement over the typically used keyword search model. The numerical pipeline using the RoBERTa architecture achieved a nMAE of 10.64% averaged across all traits.

In order to use the pipeline and the NLP models, however, a few things should be considered. As we show throughout the results, the performance of the models is dependent on several factors. First is the type of data the model is trained on, such as whether it is an expert resource, like the POWO dataset, or a more general resource like the WIKI dataset. Furthermore, we should also point out that the output of the models cannot be taken for granted. Similar to other applications of DL in ecology like animal detection in camera traps (Norouzzadeh et al., 2018), the goal is to allow for more efficient processing of large amounts of data. There is, however, a tradeoff between the amount of predicted data and precision, which the user can actively manage. The probabilistic output of the model allows users to set the threshold high in the case where they require a smaller number of predictions with high precision, or to set the threshold low to obtain a higher recall, but a lower reliability of the data requiring a longer time for manual processing. Finally, large language models can derive trait information from the contextual cues in a text (trait-trait correlations, phylogenic relationships), allowing them to potentially identify traits that might not be explicitly mentioned. While the capacity of the model to “read between the lines” can be particularly useful when dealing with ecological literature that may not always provide detailed or standardized descriptions of traits, it also means that the model can occasionally generate incorrect or unfounded predictions.

The main problem that we outline within our study is the fusion of data from two different sources. Since we used GIFT, an external database, to evaluate the traits instead of labels extracted directly from the descriptions, mismatches may arise. As a result, an optimal accuracy of 100% and a nMAE of 0% isn’t likely even in the case of a human extracting traits. For the classification examples, this is due to inconsistencies in the trait values from different sources as well as the inherent difficulty of specifying the class of some species’ traits. For example, some species like grass trees, tree ferns, semi-woody megaherbs or dwarf shrubs, bamboos, palms, climbers, etc. are difficult to squeeze into growth form categories for ecologists as well as for artificial intelligence and their treatment often differs among resources (Wenk et al., 2023). To assess this, we can use the agreement score contained in the GIFT database (Weigelt et al. 2020), representing the agreement for the trait value from different sources for each species. For the species in the POWO dataset, the average agreement score for growth form in GIFT is 92.2% with an average of 4.06 descriptions per species. In other words, if we use the primary label in GIFT to label all of the 199,241 descriptions, we get an accuracy of 92.2%. In comparison, our model has an accuracy of 92.1%, meaning that it is sufficiently high given the circumstances. For the other categorical traits, the accuracies based on the GIFT agreement scores are 99.5% for epiphyte, 93.3% for climber, 98.1% for lifecycle, and 97% for the life form. The corresponding DistilBERT accuracies were 97.9% for epiphyte, 97.1% for climber, 97.9% for lifecycle, and 86.3% for the life form. Similarly, numerical traits vary over time and space, and different floras might have different values, thus there is an inherent variation in the results reported in the literature. Furthermore, data on traits can be reported in a variety of ways. For example, the size of a plant can be reported as stem length, vegetative or reproductive height, growing height of a vine, length of a shoot of a creeping plant, and so on, making it difficult to arrive at an ultimate plant height estimate. Some resources report an average mature height while others report a rarely achieved maximum growing height. The size of leaves can be reported as lamina dimensions or blade dimensions including the petiole. In the case of compound leaves, they may refer to the leaf or leaflets. This variation in reporting makes it difficult to label even when manually processing the data (Maitner et al., 2023, Kattge et al., 2020, Wenk et al., 2023).

An additional problem of our approach is the overconfidence of the model when given small amounts of information. The training approach is based on combining independently sourced text and trait data, therefore when information for the trait is not contained in the description the model may learn erroneous relations. A way of tackling this in the future is to add negative descriptions within the training process, which contain no relevant trait information and are labeled accordingly. While the approach of fusing textual data and target descriptions from independent resources has its problems, it also has many advantages, the most important being the ease of adding a large amount of data from different resources. This allows us to easily expand the model to new traits, descriptions and even languages in a semi-automated way without the need for significant human effort involved.

There are several ways to further improve the capabilities of the model. One step is to expand the use of the model to other languages, further than using translations like in our workflow. This is important since a large number of national floras are only written in the countries’ language and this approach can expand the extent of traits that are generally biased towards the Global North and Australia. This can be done by either fine-tuning a multilingual language model such as the multilingual version of BERT or other architectures like BLOOM (Scao et al. 2022) or by fine-tuning monolingual language models like Spanish-BERT (C añete et al., 2023), GottBERT (Scheible et al., 2020) and AraBERT (Antoun et al., 2020). Another step is to adapt the domain of the transformer models (Gururangan et al., 2020) to ecological tasks, similar to what has been done in other fields such as BioBERT for Biomedicine (Lee et al., 2020) and FinBERT for Financial analysis (Liu et al., 2021). This should theoretically improve the model performance on a variety of tasks, including functional trait extraction. A foundation model for plant ecology could be used in the current pipeline, but also in tasks such as literature reviews, text summarization and named entity recognition.

Finally, the text processing pipeline can be combined with other information and data modalities. The inclusion of geographic or phylogenetic information in the form of a prior has already resulted in a large improvement in classification performance in species identification from images (Mac Aodha et al., 2019). Therefore, combining this information, as well as information such as images from the entire plant or plant parts will further increase the capabilities of the model and result in more reliable predictions.

Overall, the NLP workflow presented here holds great potential to overcome resource limitations in mobilizing plant functional trait data and may help to arrive at a more comprehensive and global understanding of functional plant ecology. While most landmark publications in functional macroecology have so far been based on trait data for up to a few thousand species including strong taxonomic and geographical biases (Maitner et al., 2023) truly global and less biased analyses seem in reach. There is a huge amount of species descriptions and trait information readily available in scientific papers, preprints, online libraries (e.g. Biodiversity Heritage Library), thematic databases (e.g. JSTOR Global Plants), university library digitization programs (e.g. MINE, BIOfid), monographs and floras (Frodin, 2001), and public websites (e.g. Wikispecies). We argue that in combination with targeted field campaigns in undersampled regions, the mobilization of already available but unstructured information in texts from the above resources may help to fill gaps in pertinent trait databases and hence boost the availability of trait data for macroecological analyses.

## Author Contributions

Viktor Domazetoski, Patrick Weigelt and Holger Kreft conceived the ideas of the paper. Viktor Domazetoski and Patrick Weigelt designed the methodology alongside helpful discussions with Philipp Wieder, Radoslav Koynov, Alireza Zarei and Holger Kreft. Viktor Domazetoski performed the analyses and created the visualizations. Viktor Domazetoski led the writing of the first draft of the manuscript. All authors contributed critically to the drafts and gave final approval for publication.

## Acknowledgements

HK acknowledges funding from the DFG as part of the research unit FOR 2716 DynaCom.

## Footnotes

1 The code to train, evaluate and use the models is available at https://github.com/ViktorDomazetoski/NLP-Plant-Traits

