## Supplementary Information S1 for "Using natural language processing to extract plant functional traits from unstructured text"

### S1. Pipeline Example

Figure S1: Example application of the proposed NLP pipeline used for the prediction of

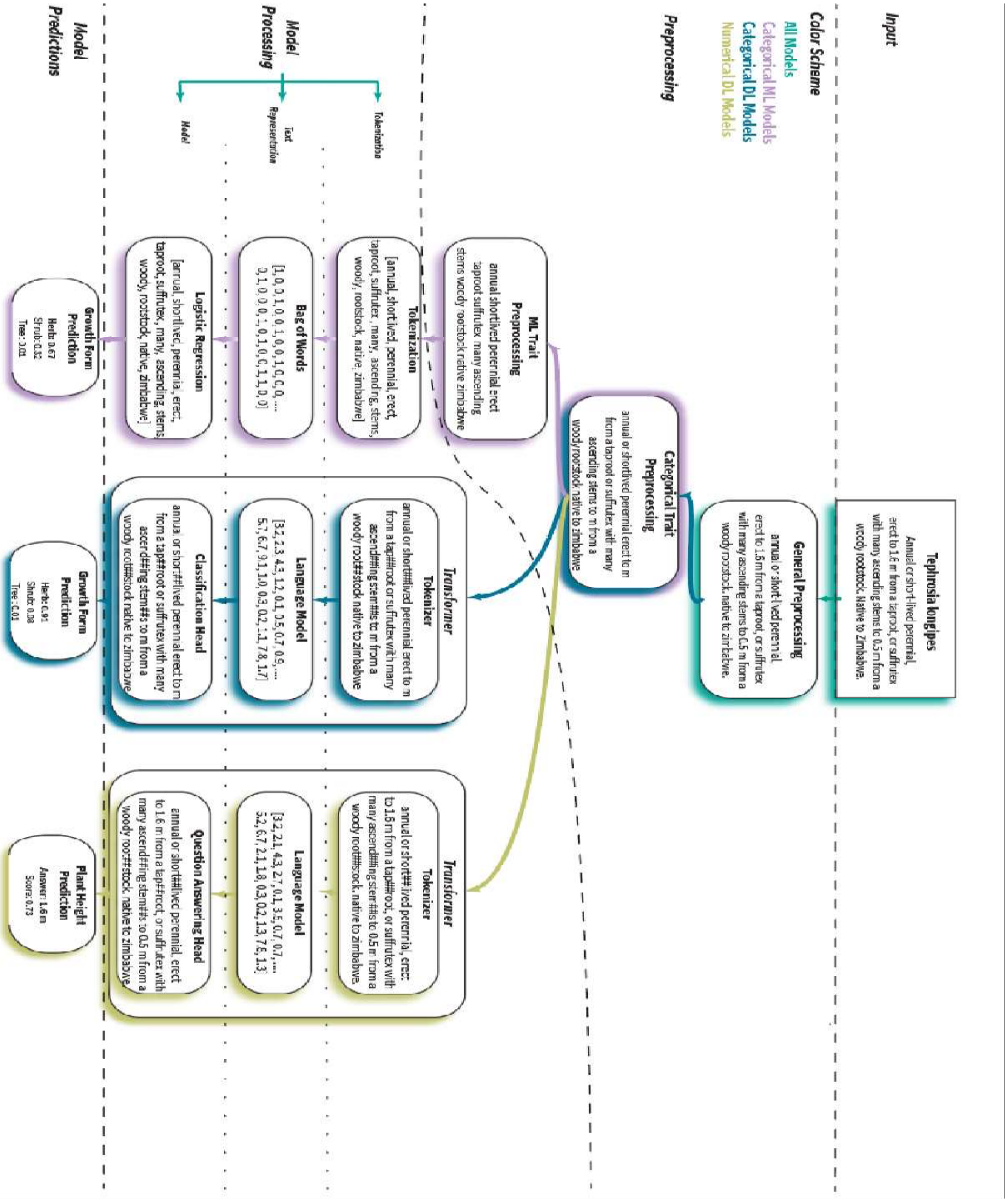


categorical and numerical traits for one species. The description first goes through a general preprocessing pipeline where it is lowercased and all abnormalities are removed. Then for the categorical traits, the digits and punctuation are further removed and only for the categorical ML pipeline, stop words are also removed. The preprocessed description is then tokenized based on the requirements of the model and is embedded in a vector space using the bag of words model or a large language model. Using this embedding as input for a classification or question answering model, the model associates weights to each trait value (only one shown here). Using these weights, the final prediction is made and an associated confidence score is provided.
