## Supplementary Information S2 for "Using natural language processing to extract plant functional traits from unstructured text"

### S2.1 Plants of the World Online Dataset

Plants of the World Online (http://www.plantsoftheworldonline.org/) is an online platform established by Kew Royal Botanic Gardens, dedicated to digitizing and sharing comprehensive data on the world's flora. The Plants of the World Online scrape comprised 288,254 descriptions encompassing 59,151 species, categorized into 251 distinct description types, originating from 15,023 different sources. On average, each species is associated with approximately 4.86 descriptions (standard deviation 5.84), though due to an uneven distribution, the median count is 2 descriptions per species. To create the POWO dataset, these descriptions were organized on a per-species-per-source basis. On average, the descriptions in this aggregated POWO dataset consist of 118 words, however, the median is only 6 words (Fig. S2a). This discrepancy is primarily attributed to the fact that a significant portion (46.9%) of the descriptions contain fewer than five words, necessitating manual examination to determine if they contain the trait data required for training and evaluating models. The entire corpus of descriptions comprises 52,336 tokens, with 44% occurring only once and 80% occurring fewer than ten times (Fig. S2b) The geographic species distribution of the dataset closely mirrors the actual distribution of plant species (Fig S3a). However, the average geographic word count reveals variation, with the Amazon region, having the highest number of species descriptions, featuring the fewest words, averaging under 50 words per species (Fig S4a). In contrast, descriptions in Africa are much more extensive, as the initial purpose of Plants of the World Online was to document these species. In terms of trait coverage, the growth form, epiphyte and climber traits have a substantial representation, surpassing 70%, while numerical traits, such as leaf length and width, have a much more limited coverage, with only a few thousand descriptions available (Table S1).

Table S1: Trait and trait value coverage for the POWO dataset.

| Dataset | Trait Name | Trait Count (#) | Trait Coverage (%) | | |
|---|---|---|---|---|---|
| **Categorical Traits** | | | | **Trait Class** | **Trait Class Coverage (%)** |
| POWO | Growth Form | 46110 | 78 | Herb | 59.3 |
| | | | | Shrub | 17.1 |
| | | | | Tree | 23.5 |
| POWO | Epiphyte_1 | 44417 | 75.1 | Epiphyte | 9.8 |
| | | | | Terrestrial | 89.3 |
| POWO | Climber_1 | 45608 | 77.1 | Climber | 9.5 |
| | | | | Self-supporting | 90.2 |
| POWO | Lifecycle_1 | 36361 | 61.5 | Annual | 6.8 |
| | | | | Perennial | 93.1 |
| POWO | Life_form_1 | 21400 | 36.2 | Phanerophyte | 27.7 |
| | | | | Chamaephyte | 8.5 |
| | | | | Hemicryptophyte | 33.6 |
| | | | | Cryptophyte | 10.7 |
| | | | | Therophyte | 19.5 |
| **Numerical Traits** | | | | **Trait Mean** | **Trait Standard Deviation** |
| POWO | Plant_height_max | 17648 | 29.8 | 7.42 m | 10.67 m |
| POWO | Leaf_length_max | 3397 | 5.7 | 12.87 cm | 31.06 cm |
| POWO | Leaf_width_max | 2243 | 3.8 | 6.54 cm | 95.43 cm |

### S2.2 Wikipedia Dataset

The English Wikipedia Corpus scrape comprises 194,994 descriptions related to 55,631 species, organized into 7,903 description types, sourced from 22,035 unique authors. On average, there are about 3.5 descriptions for each species, with a standard deviation of 3.1. However, due to a skewed distribution, the median number of descriptions per species stands at 3. These descriptions were consolidated into a single description per species per source, resulting in the formation of the WIKI dataset, encompassing 55,631 unique entries. The descriptions of the WIKI dataset exhibit an average of 198 words, with a median length of 98 words (Fig. S2a). In total, the corpus of descriptions consists of 211,645 tokens, with 48\% of them occurring only once and 84\% occurring less than ten times (Fig. S2b). Many of these tokens with few occurrences contain taxonomic information about the species, such as family, genus or binomial names. The geographic distribution of species in the dataset exhibits a bias towards English-speaking countries, aligning with the expectations from the English Wikipedia Corpus (Fig. S3b). The average word count displays a more even distribution globally (Fig. S4b). Nevertheless, some regions, such as the Amazon, remain relatively unrepresented in the dataset.

This observation suggests that sourcing data from Wikipedia in various languages could potentially augment the dataset with new species and information. Trait coverage distribution in the WIKI dataset is similar to the POWO dataset, with generally higher coverage, as the data scraping specifically targeted species with corresponding label data in GIFT. (Table S2).

Table S2: Trait and trait value coverage for the WIKI dataset.

| Dataset | Trait Name | Trait Count (#) | Trait Coverage (%) | | |
|---|---|---|---|---|---|
| Categorical Traits | | | | Trait Class | Trait Class Coverage (%) |
| WIKI | Growth Form | 49535 | 89 | Herb | 49.8 |
| | | | | Shrub | 20.8 |
| | | | | Tree | 29.3 |
| WIKI | Epiphyte_1 | 45022 | 80.9 | Epiphyte | 11.1 |
| | | | | Terrestrial | 88.2 |
| WIKI | Climber_1 | 50517 | 90.8 | Climber | 3.5 |
| | | | | Self-supporting | 96.3 |
| WIKI | Lifecycle_1 | 44569 | 80.1 | Annual | 7.6 |
| | | | | Perennial | 92 |
| WIKI | Life_form_1 | 25048 | 45 | Phanerophyte | 39.1 |
| | | | | Chamaephyte | 6.8 |
| | | | | Hemicryptophyte | 16.6 |
| | | | | Cryptophyte | 19.3 |
| | | | | Therophyte | 18.3 |
| Numerical Traits | | | | Trait Mean | Trait Standard Deviation |
| WIKI | Plant_height_max | 23696 | 42.6 | 6.67 m | 10.82 m |
| WIKI | Leaf_length_max | 8159 | 14.7 | 16.1 cm | 103.87 cm |
| WIKI | Leaf_width_max | 7116 | 12.8 | 4.92 cm | 53.87 cm |

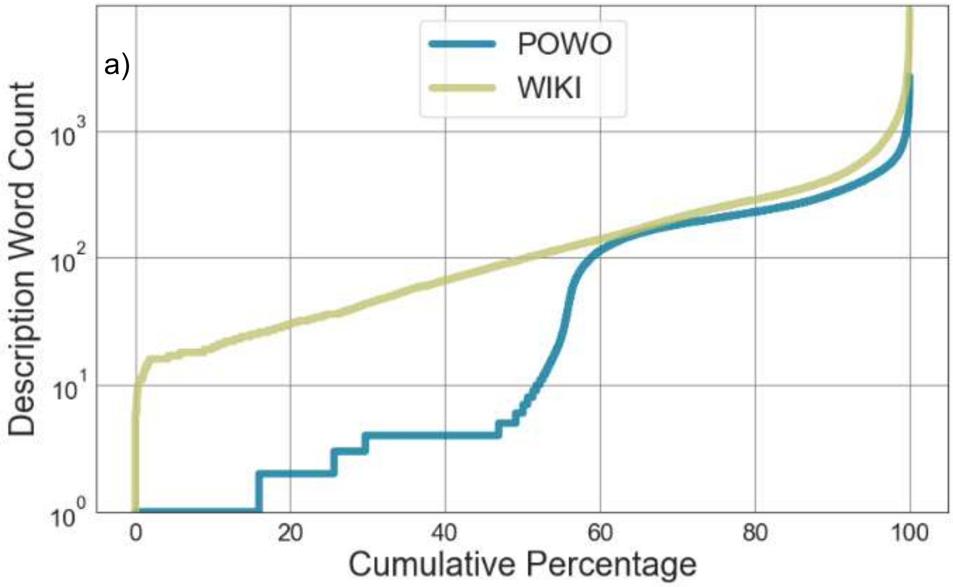


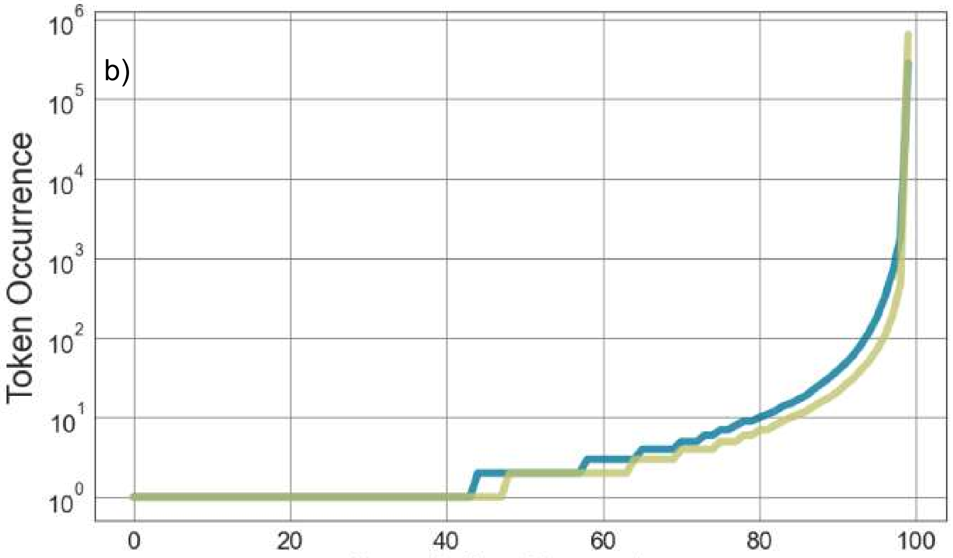


Figure S2: Cumulative percentage of the log-description word count (a) and log-token occurrence (b) of the Plants of the World Online (POWO) (blue) and Wikipedia (WIKI) (olive).

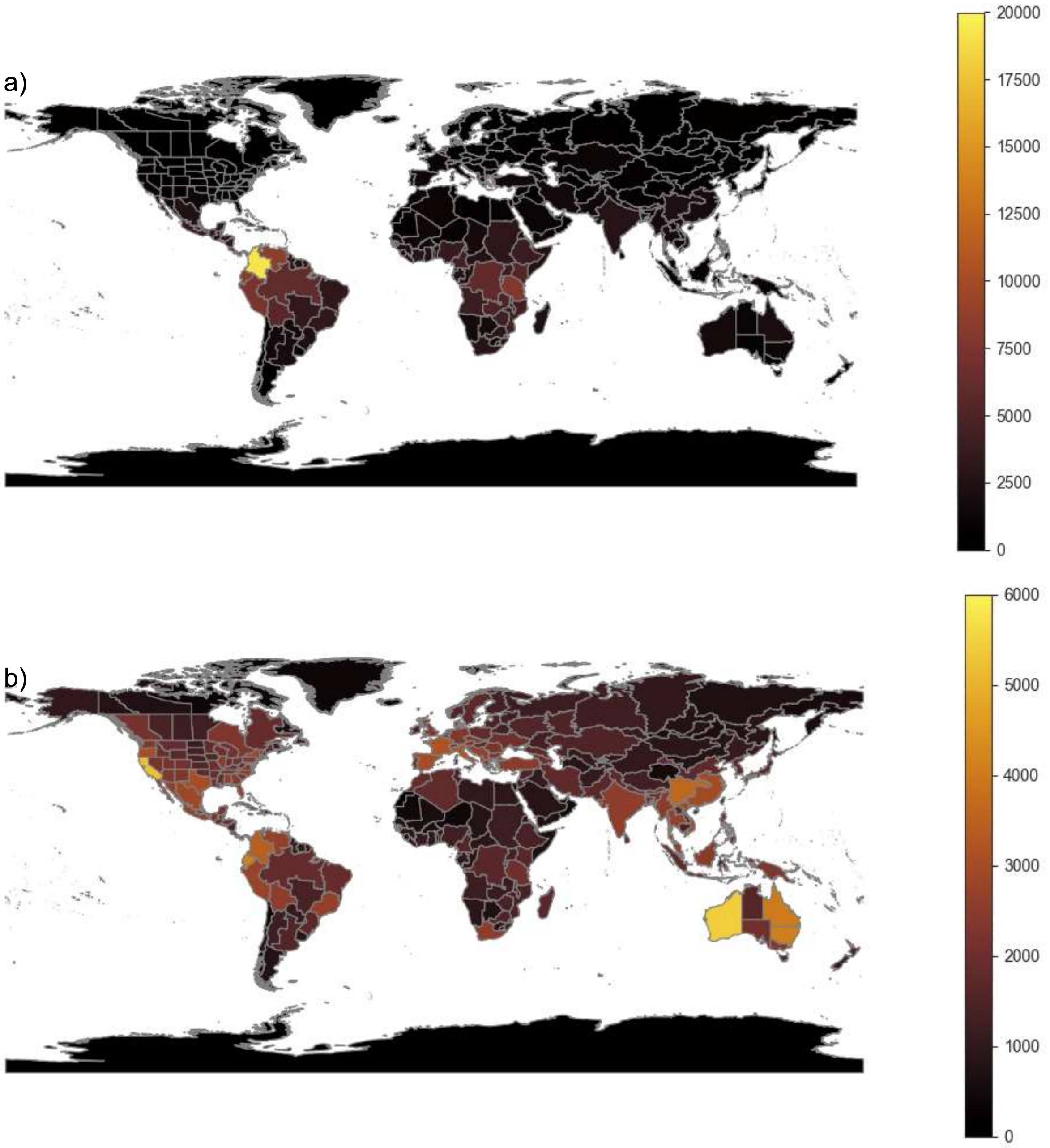


Figure S3: Geographic distribution of the species in the Plants of the World Online (POWO) (a) and Wikipedia (WIKI) (b) datasets. To map the species distribution, we combined the 59,151 species of the POWO and 55,631 species of the WIKI dataset with their level 3 botanical region of the World Checklist of Vascular Plants (Govaerts et al., 2021) and then calculated the species richness for each botanical region.

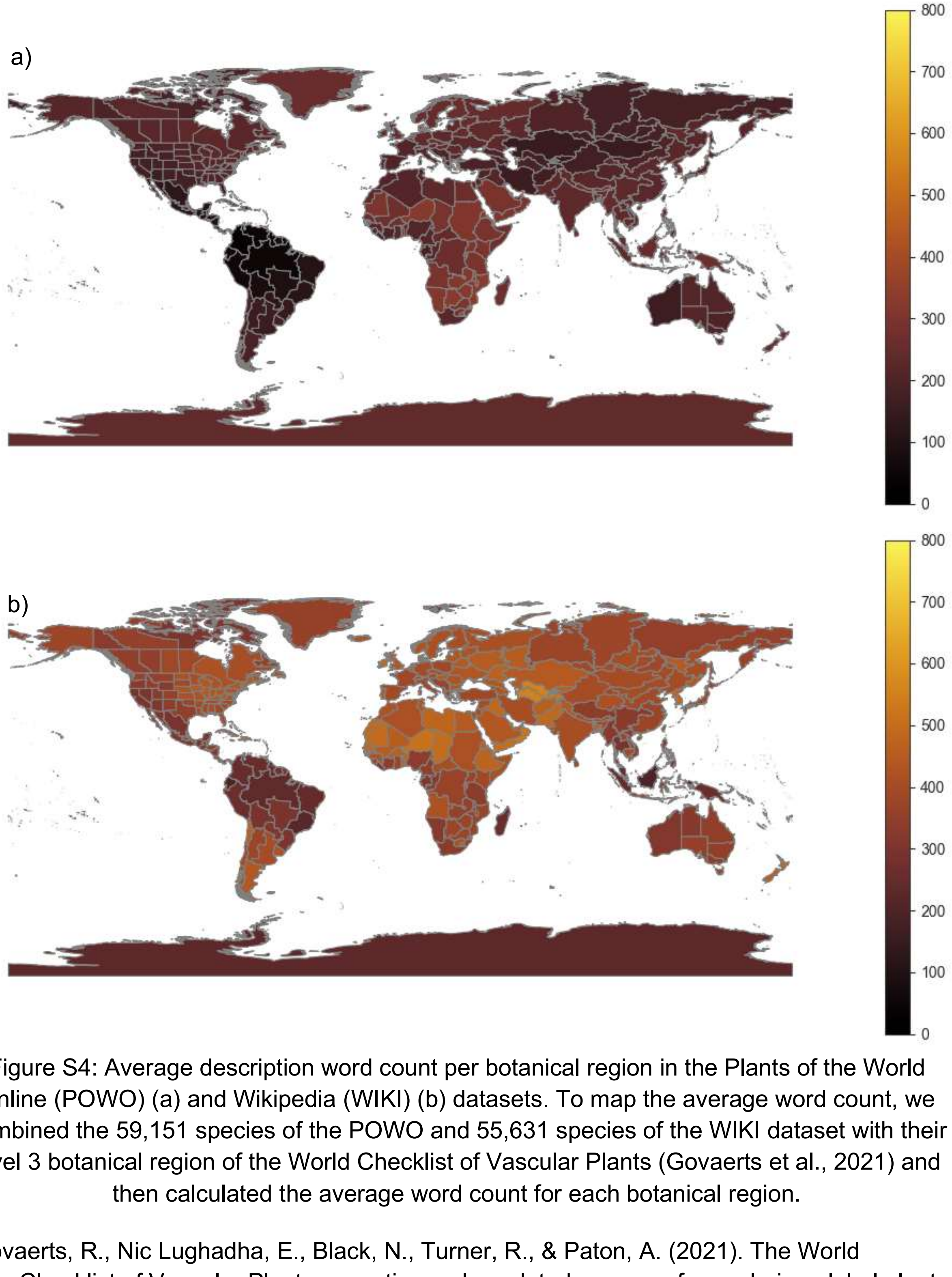


Figure S4: Average description word count per botanical region in the Plants of the World Online (POWO) (a) and Wikipedia (WIKI) (b) datasets. To map the average word count, we combined the 59,151 species of the POWO and 55,631 species of the WIKI dataset with their level 3 botanical region of the World Checklist of Vascular Plants (Govaerts et al., 2021) and then calculated the average word count for each botanical region.

Govaerts, R., Nic Lughadha, E., Black, N., Turner, R., & Paton, A. (2021). The World Checklist of Vascular Plants, a continuously updated resource for exploring global plant diversity. Scientific Data, 8(1), 215.
