## Supplementary Information S3 for "Using natural language processing to extract plant functional traits from unstructured text"

### S3. Regex Keywords

| | | |
|---|---|---|
| **Growth Form** | Herb | herb, herbaceous, forb, orchid |
| | Shrub | shrub, subshrub, undershrub, bush, shrublet, cactus |
| | Tree | tree, mallet |
| **Epiphyte** | Epiphytic | epiphyte, epiphytic |
| | Terrestrial | terrestrial |
| **Climber** | Climber | climber, vine, liana, liane, woodytwiner, shrubbytwiner, woodyclimber, twiner |
| | Self-supporting | self-supporting |
| **Lifecycle** | Annual | annual |
| | Perennial | perennial |
| **Life Form** | Phanerophyte | phanerophyte |
| | Chamaephyte | chamaephyte |
| | Hemicryptophyte | hemicryptophyte |
| | Cryptophyte | cryptophyte |
| | Therophyte | therophyte |
