## Supplementary Information S4 for "Using natural language processing to extract plant functional traits from unstructured text"

### S4. Model Implementation

The codebase was written in Python v. 3.9.13 and either directly or indirectly relied heavily on the NumPy (Harris et al., 2020) and pandas (McKinney, 2021) libraries for the organization and processing of data. Text analysis and preprocessing were done using the NLTK python library (Loper & Bird, 2002). We implemented the BOW and logistic regression model using the scikit-learn ML library v. 1.1.3 (Pedregosa et al., 2011). We trained and evaluated the large language models using the Huggingface's transformers library v. 4.28.0 (Wolf et al., 2019). The models were trained using Google Colab and Kaggle on the freely available T4 GPUs. The models were trained for 3 epochs with a batch size of 16 and a maximum sequence length of 512 tokens. The default learning rate was set to $2e^{-5}$ and a weight decay of 0.01 was applied. The visualizations were done using the Python Matplotlib (Hunter, 2007) and Seaborn libraries (Waskom, 2021). The entire codebase for the manuscript is open-source and available on https://github.com/ViktorDomazetoski/NLP-Plant-Traits.


Harris, C. R., Millman, K. J., Van Der Walt, S. J., Gommers, R., Virtanen, P., Cournapeau, D., ... & Oliphant, T. E. (2020). Array programming with NumPy. Nature, 585(7825), 357-362.

McKinney, W. (2011). pandas: a foundational Python library for data analysis and statistics. Python for high performance and scientific computing, 14(9), 1-9.

Pedregosa, F., Varoquaux, G., Gramfort, A., Michel, V., Thirion, B., Grisel, O., ... & Duchesnay, É. (2011). Scikit-learn: Machine learning in Python. the Journal of machine Learning research, 12, 2825-2830.

Wolf, T., Debut, L., Sanh, V., Chaumond, J., Delangue, C., Moi, A., ... & Rush, A. M. (2019). Huggingface's transformers: State-of-the-art natural language processing. arXiv preprint arXiv:1910.03771.

Hunter, J. D. (2007). Matplotlib: A 2D graphics environment. Computing in science & engineering, 9(03), 90-95.

Waskom, M. L. (2021). Seaborn: statistical data visualization. Journal of Open Source Software, 6(60), 3021.

Loper, E., & Bird, S. (2002). Nltk: The natural language toolkit. arXiv preprint cs/0205028.
