## Supplementary Information S5 for "Using natural language processing to extract plant functional traits from unstructured text"

### S5. Inter-dataset evaluation

In order to effectively utilize these models in practical applications, it’s crucial to assess their performance on datasets with distinct characteristics from those used for training. To address this, we conducted an inter-dataset evaluation, where models trained on the training set of one dataset were evaluated on the test set of the remaining two datasets (Fig. S5). Along the diagonal lie the intra-datasets scores. The results revealed that the models trained on POWO and POWO_MGH demonstrated nearly identical intra-dataset performance. This implies that, particularly for categorical models, utilizing only the POWO categories directly relevant to a specific trait does not significantly enhance model performance. However, in the case of inter-dataset scores, the variation in performance between datasets remained relatively consistent across traits, with one notable exception: the life form trait. The observed disparity can be attributed to the limited data available in the POWO_MGH dataset for this specific trait. When evaluating inter-dataset scores between POWO and WIKI, a more pronounced decline in performance was evident across all traits. This decrease in performance averaged 8.9% in precision and 23% in recall when POWO served as the training dataset, and 13.8% in precision and 8.5% in recall when WIKI was the training dataset. The most substantial decrease was again observed in the life form trait, with precision decreasing by 25% (14%) and recall by 26% (13%) for the POWO (WIKI) datasets. These findings underscore the importance of exercising caution when drawing inferences from new data, as the model's performance is intricately tied to the characteristics of the training dataset.

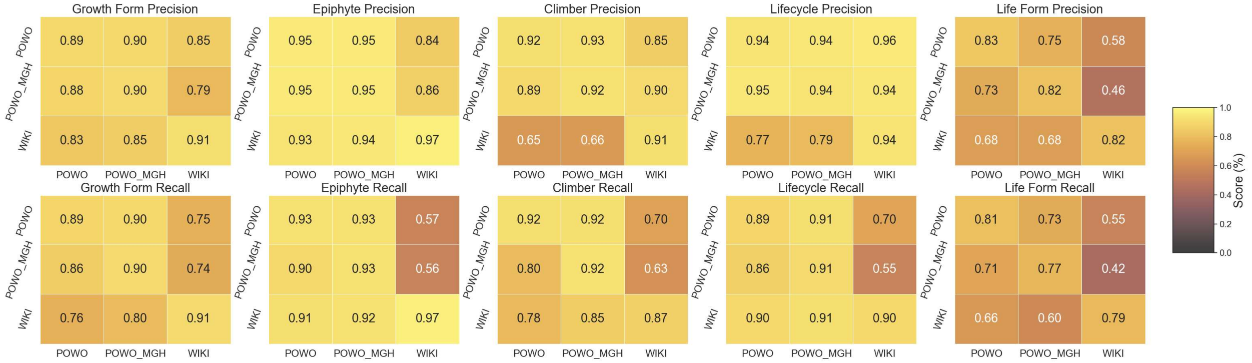


Fig. S5. Inter-dataset precision (first row) and recall (second row) scores using the DistilBERT model for the categorical traits of interest (columns). Within each matrix, each row of the heatmap represents the dataset used in training, while each column represents the dataset used for testing. The values on the diagonal correspond to the values of the intra-dataset models discussed in section 3.1.1.
