## Supplementary Information S6 for "Using natural language processing to extract plant functional traits from unstructured text"

### S6. Analysis of probabilistic predictions

We also explored the impact of the probability threshold (t) on model performance. This threshold serves as a versatile tool, enabling users to tailor results according to their specific requirements. Raising the threshold allows users to achieve a higher precision at the expense of a reduced recall while lowering the threshold yields the opposite effect. To gain insights into this behavior, we constructed precision-recall curves. These curves provide a visual representation of how the model's performance evolves and how the trade-off between precision and recall varies at different threshold settings (see Fig. S6).

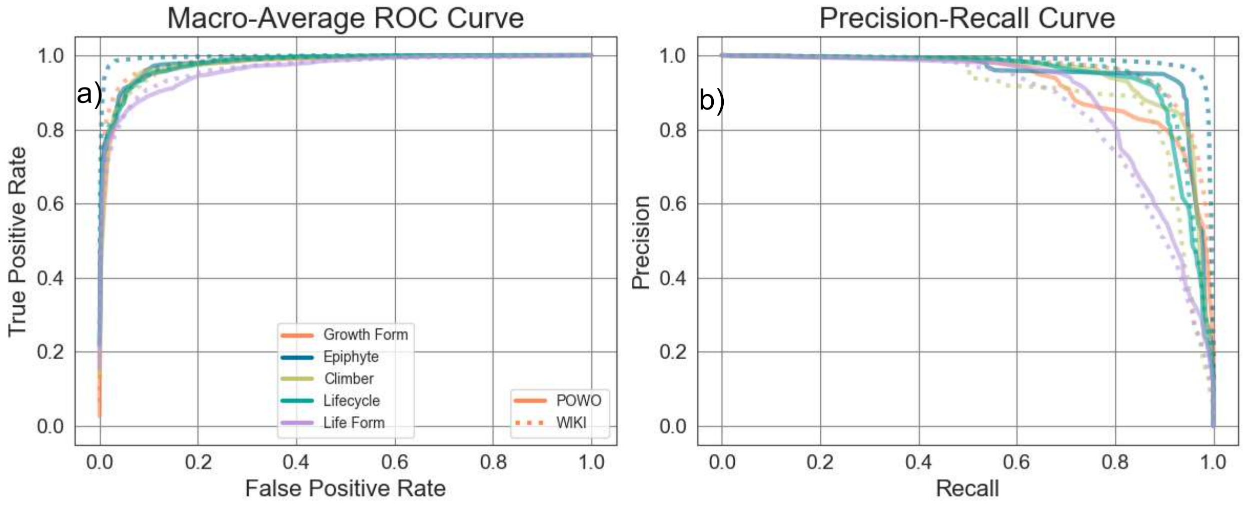


Fig. S6. Macro-average ROC curve (a) and Precision-Recall curve (b) for the DistilBERT model on the Plants of the World Online (POWO) (solid line) and Wikipedia (WIKI) (dotted line) datasets.

For a more comprehensive understanding of how specific threshold values affect the models, we generated precision and recall curves across a range of threshold values from 0 and 1 (Fig. S4). As anticipated, precision displayed a positive correlation with the threshold, peaking at optimal values around t=0.8. However, for select traits (epiphyte, climber, lifecycle), the precision curve plateaued at around t=0.4, while for the growth form and life form traits, precision continued to rise until the higher end of the threshold range. Conversely, recall exhibited the opposite pattern, with a more consistent performance across the five traits of interest.

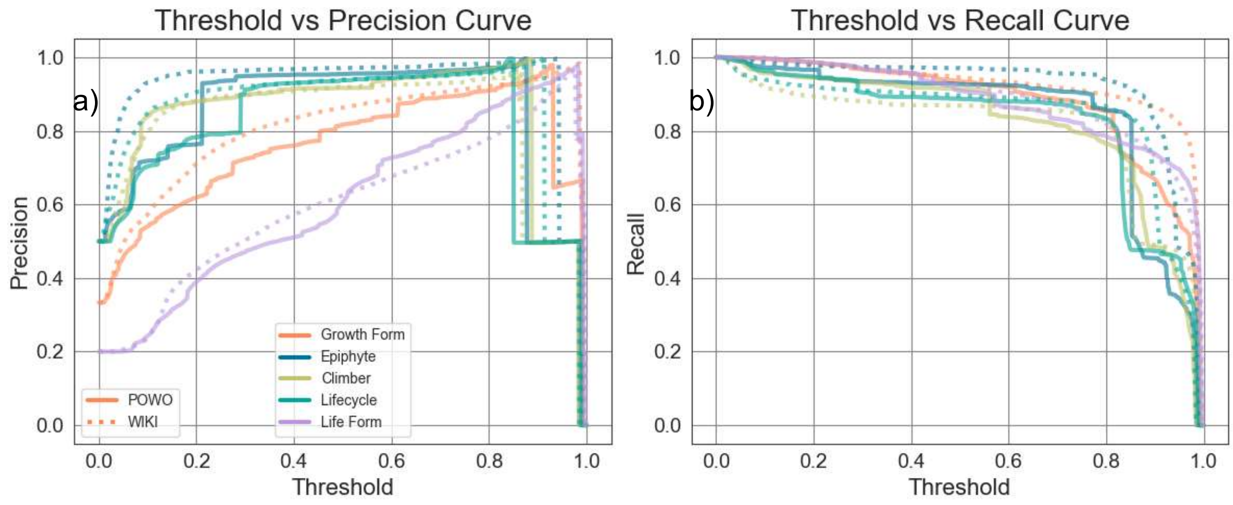


Fig. S7. Threshold vs Precision (a) and Recall (b) for the DistilBERT model on the Plants of the World Online (POWO) (solid line) and Wikipedia (WIKI) (dotted line) datasets.

Finally, we contrast the results reported in the main section of the manuscript, which were obtained by taking the trait prediction with the highest probability (argMax), with the model's performance when applying probabilistic thresholds of *t=0.5* and *t=0.8* (Fig. S8). We observe that when using *t=0.5,* the results generally resemble those of the argMax approach, albeit sometimes at the expense of precision in favor of achieving higher recall, such as in the growth form and life form traits. On the other hand, t=0.8 yields a substantial increase in precision, at the cost of a reduced recall, making it valuable for scenarios where the data quality must be stringent and manual verifications are limited.

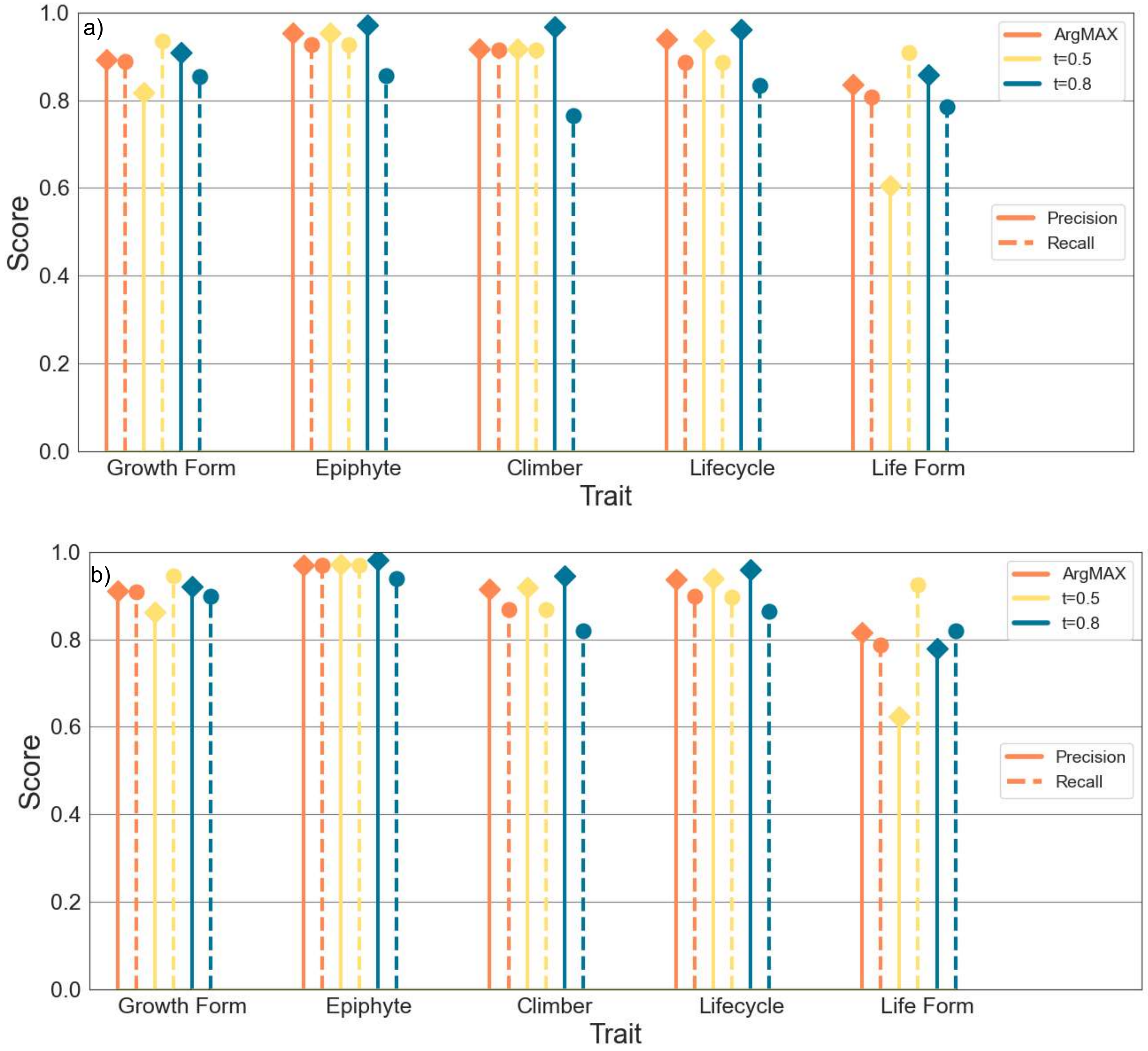


Fig. S8. Precision (solid line) and Recall (dashed line) for the DistilBERT model on the Plants of the World Online (POWO) (a) and Wikipedia (WIKI) (b) datasets when using an argMax approach (orange), a threshold t=0.5 (yellow) and t=0.8 (blue).
